# Investigating graph neural network for RNA structural embedding

**DOI:** 10.1101/2022.12.02.515916

**Authors:** Vaitea Opuu, Hélène Bret

## Abstract

The biological function of natural non-coding RNAs (ncRNA) is tightly bound to their molecular structure. Sequence analyses such as multiple sequence alignments (MSA) are the bread and butter of bio-molecules functional analysis; however, analyzing sequence and structure simultaneously is a difficult task. In this work, we propose CARNAGE (Clustering/Alignment of RNA with Graph-network Embedding), which leverages a graph neural network encoder to imprint structural information into a sequence-like embedding; therefore, downstream sequence analyses now account implicitly for structural constraints. In contrast to the traditional “supervised” alignment approaches, we trained our network on a masking problem, independent from the alignment or clustering problem. Our method is very versatile and has shown good performances in 1) designing RNAs sequences, 2) clustering sequences, and 3) aligning multiple sequences only using the simplest Needleman and Wunsch’s algorithm. Not only can this approach be readily extended to RNA tridimensional structures, but it can also be applied to proteins.

## 1 Introduction

For only a few decades, the discovery of natural enzymatic RNA (ribozymes) made the search and analysis of non-coding RNA crucial in many fields of molecular biology. Alignment of RNA sequences of nucleotides is the approach of choice for such a problem; however, the RNA molecular structures are more conserved over the set of molecules sharing the same biological function than the sequences of nucleotides. By taking into account explicitly the molecular structures of RNAs, specialized alignment algorithms allow high-quality alignments compared with structure-free methods (19).

In contrast with proteins, RNA molecular structures can be simplified to their so-called secondary structure where only canonical base pairs (BP) such as G-C, A-U, and G-U are considered. The minimum free energy (MFE) secondary structure of an RNA molecule can be efficiently determined by combining a model of physical interactions devised by Mathew and coworkers (10) and the Zucker algorithm (21) to identify the minimum free energy structure (MFE). The opposite approaches use various supervised learning methodologies such as covariance models (2) or deep neural networks (14). However, recent work from (3) revealed that neural network-based methods suffer from generalization issues, making these supervised methods less suitable for discovering new motifs or functions.

The alignment problem is often a supervised task in which a collection of (handcrafted) multiple sequence alignments (MSA) are used to build scoring systems called substitution matrices, *e.g*. BLOSUM62 (6). The modern era using deep learning is no exception. For RNA molecules, a recently proposed method using BERT-like models (1) trained on sequence masking and explicit structural alignments gave very high performances in alignment benchmarks. Before this, SAdLSA (4) was introduced to protein alignments with the same approach but using CNN. These approaches were introduced such that the molecular structure constraints are accounted for implicitly while aligning sequences using Needleman and Wunsch’s algorithm (12).

We propose here a trained projection called CARNAGE that creates a vector representation of RNA sequences that implicitly incorporate the molecular structure. This encoding is then used in downstream analyses such as sequence alignment, clustering, or motif searching that now implicitly account for structure. Our approach leverages the graph neural network architecture as shown in Fig 1.

**Figure 1:**
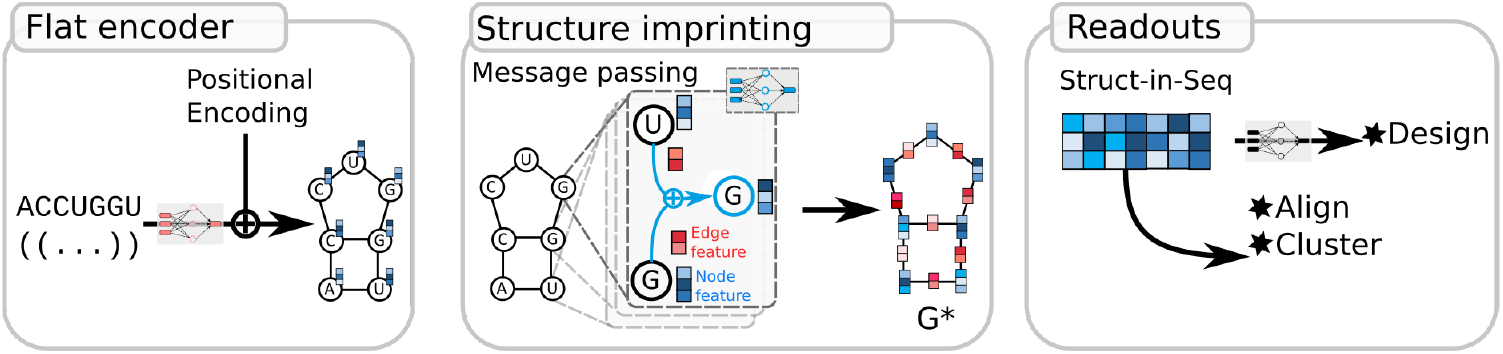
**The CARNAGE framework for sequence embedding. Flat encoder module**: a neural network is first applied to nucleotides, with which the positional encoding is summed. Then, the sequence is turned into a graph using the molecular structure. **Structure imprinting**: for each node/nucleotide, two rounds of message passing network aggregate information from the neighbors. The output is a collection of vectors representing the sequence. The node vector representation now contains structural information. **Readouts**: all the node vectors are concatenated to form the Si-seq, which is used for design, alignment, or clustering.

To assess the benefits and inconveniences of our approach, we conducted two types of investigations:

- Synthetic data: based on synthetic data – random sequences or structures— we showed that our embedding not only encodes some sequence and structural information but can also be successfully used in comparing sequences.
- Real data: we showed that our approach, although unsupervised, is comparable to a well-known method either based only on sequences or based on structure-folding.

## 2 Methods

### 2.1 Neural network architecture

The input of our model is a sequence and its predicted structure. We first create a graph *G* = (*V, E, U*), where nodes *V* are unit-vectors encoding the nucleotide identity, for example Adenine is encoded with (1,0,0,0); E are edges connecting adjacent nucleotides and base-paired positions; and U is a vector that encodes information about the graph G. In the beginning, only the nodes have features.

A first encoding linear block is applied to each node in order to enrich its representation then the same positional encoding (PE) used in the successful Transformer model (16) is summed to it (see Fig. 1). Next, we perform two encoding rounds of message passing (MP). Each MP is composed of two updating schemes:

- The edge features are updated using the adjacent nodes and their current state. In the first round, it only uses the adjacent node features.

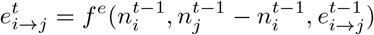
- Each node’s features are updated by aggregating “messages” from *i* neighbors 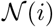.

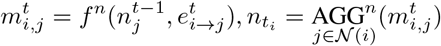
- the global node is then updated by aggregating messages from all nodes.

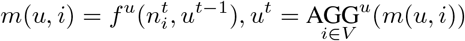

*f^e^, f^n^*, and *f^u^* are neural networks whereas AGG^*n*^, and AGG^*u*^ are respectively the sum and average aggregating functions. After the two rounds of MP, we obtain a graph *G*\* (see Fig. 1) where the features of edges ∈ ℝ^16^, nodes ∈ ℝ^64^, and the global node ∈ ℝ^64^ are richer. The final set of node features n* are concatenated to form the sequence embedding we call Si-seqs.

The input data for this training is a collection of pairs (sequences, structure). First, for each sequence-structure, we mask the identity of 5% of the positions by putting all the components of the input unit vector to zero. Then, the MP is applied to create Si-seqs. Second, we apply a final network called readout on each node features to predict the likelihood of all 4 nucleotides at each masked position:

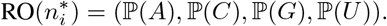

A cross-entropy loss is used to optimize parameters.

For this work, we trained the parameters on synthetic data: the training is done on random sequences and their predicted structures (obtained with ViennaRNA package (9)) with lengths *l ~ U* (50,150). We performed 500 optimization steps with the RMSprop algorithm. The learning rate is set to *lr* = 10^-3^ for batches of 64 (sequence, structure) pairs. One consequence of this strategy is that training should not be affected by finite-size dataset effects.

### 2.2 Alignment score

We used Needleman & Wunsch’s alignment algorithm, where we only replaced the similarity score. The traditional pairwise alignment algorithm starts with two sequences, X and Y of length *l_X_, l_Y_*, where the dynamic programming matrix *F* is filled with the recursion:

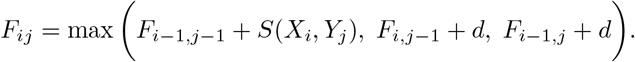

*S*(*X_i_,Y_j_*) is traditionally the similarity between nucleotides *X* and *Y* at positions *i* and *j* respectively. *d* is the penalty score set for the experiments to *d* = −4. The alignment score is *F_l_x_, l_Y__*.

Now, we replaced the similarity of nucleotides with the euclidean distance between the nucleotides embedding 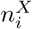 and 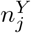:

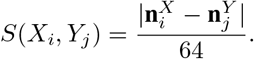

For the case of multiple sequence alignments, we start by computing the alignment scores of all pairs of sequences. We then create the guide tree for the alignment using a hierarchical clustering method implemented in scipy (17). By following the guide tree, we iteratively make pairwise alignments. Once a pairwise alignment is formed, the embeddings of the pairs are combined by averaging over the embedding components.

## 3 Results on synthetic data

To what extent the structure encoded in si-seqs correlates to the actual structure? We crafted a dataset of 30 sequences designed to fold into structure A and 30 others designed to fold into different structure B (see Fig. 2). We chose to design the sequences using a simple genetic algorithm optimization. For each pair of sequences, we computed an alignment score of Si-seqs. The pairwise scores obtained were next fed to a hierarchical clustering algorithm implemented in `scipy`. Fig 2 shows the dendrogram obtained where two groups have been separated: the sequences folding in structure A and the sequences folding in structure B.

**Figure 2:**
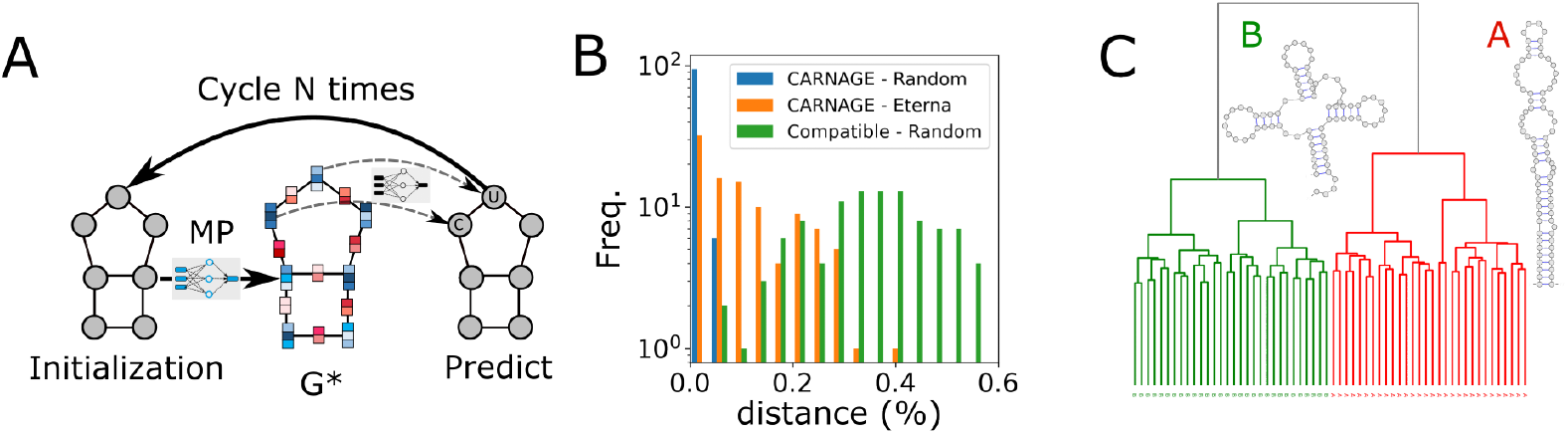
**A** Design RNA sequences. We randomly initialize the graph nodes; then, we apply the MP to obtain the node embedding for all positions and edges. Next, 5% of the positions are assigned nucleotide probabilities using the readout network (see above). The updated graph is fed again to the MP. **B** Histogram of the design quality for random and Eterna structures measured with the hamming distance between the targeted structures and the predicted structure of the designs. In green, we show the score of sequences sampled with the only constraint of having canonical BPs at paired positions. **C** Clustering of synthetic sequences. Dendrogram of 30 sequences designed for structures A and 30 others for structures B. The distance matrix was computed with Si-seqs alignment scores.

To design sequences, we start by creating a graph where the connections are given by the targeted structure. In the case of design, the nodes are yet to be determined. As shown in Fig 2, we randomly initialize all the nodes’ features using the uniform distribution. Then, we applied multiple rounds of MP to create the Si-seq. Next, we randomly pick 5% of the positions in the initial graph and replace them with the predicted likelihood 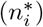. We applied this encoding-prediction step iteratively for 200 distance (%) steps. Finally, we extracted the most likely nucleotide type per position as the predicted sequence. To assess the quality of the design, we used the hamming distance:

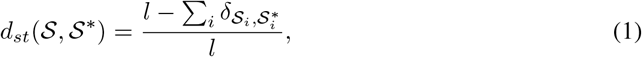

where 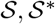 are the predicted and targeted structures, respectively in the dot-bracket notation. 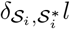 is Kronecker delta which equals 1 if 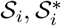 are the same symbol.

We applied the design procedure to 100 random structures obtained by sampling random sequences of length 50 ≤ *l* ≤ 150. We performed 10 design runs per structure. 70 out of 100 designed sequences fold exactly in their targeted structure. On the RNA designed benchmark Eterna (8), only 18 fold exactly in their targeted structures. Finally, we performed for each random target structure a sampling of 1000 compatible sequences *i.e*. sequences where base-paired positions follow the canonical pairs of bases {GC, GU, AU}; and selected the best one in terms of structure distance. This last test showed that the design task could not be trivially solved by respecting the canonical pairs of bases; therefore, our method learns more than canonical pairs. Fig 2 recapitulates the design performances.

## 4 Results on natural data

We assessed first the quality of our method on the sequence clustering task. We extracted randomly 23 families from RFAM (5) with average sequence length l within 80 ≤ l ≤ 150. For the sake of representation, we only selected 10 sequences per family (230 sequences in total). Fig 3 shows the dendrogram obtained from the pairwise alignment scores where the 230 sequences have been grouped into 27 clusters using the hierarchical clustering algorithm minimizing squared distances within clusters (implemented in `sklearn` (13)). We applied the same procedure with two other programs: i) `MAFFT` (7) that uses only the sequence to create the MSA and ii) `Locarna` (18) that uses both the structure and the sequence. Our results showed that `Locarna` (perfect clustering) and `MAFFT` performed better than our approach. Only 22 pure clusters were found, where 16 are composed of 10/10 and 9/10 sequences of the same family. The obtained clusters displayed 0.92 for homogeneity, 0.95 for completeness, and 0.93 for V-measure (the arithmetic mean of homogeneity and completeness).

**Figure 3:**
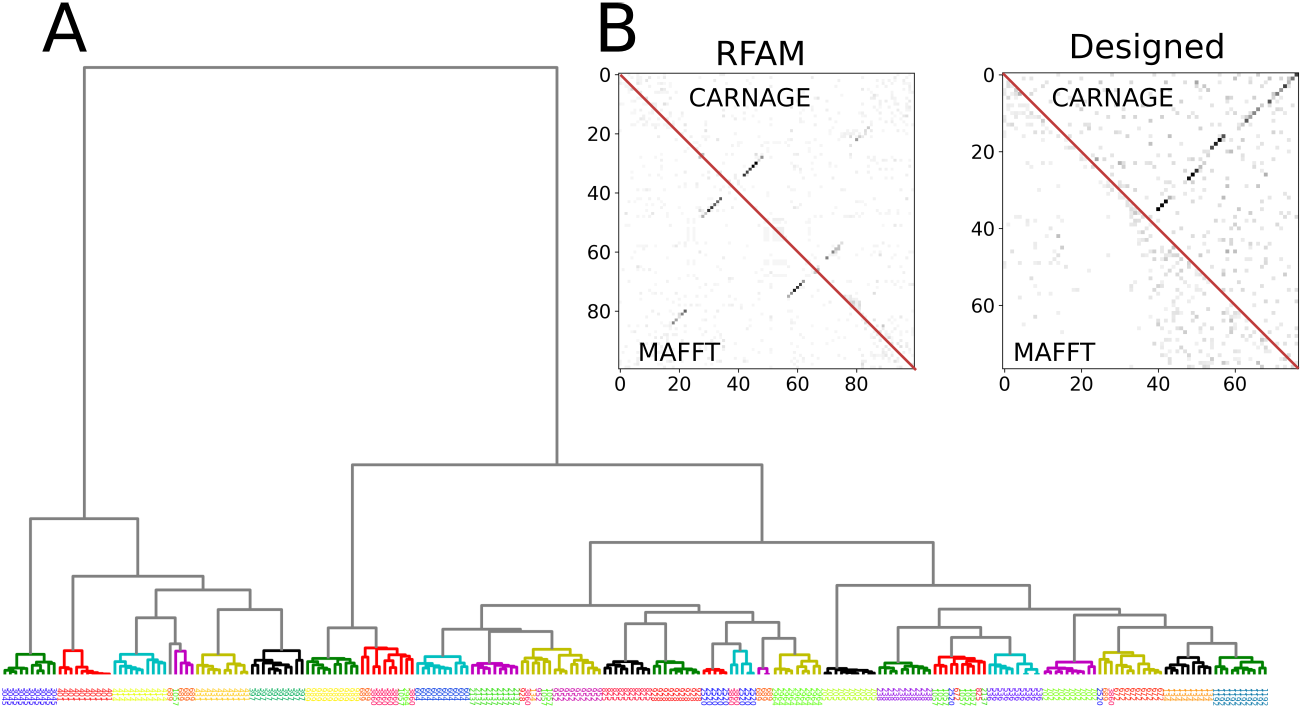
Application to real data. A) A second commonly used analysis is coevolution analysis. A Dendrogram where the formed clusters are colored on the tree, whereas the labels are colored by RFAM family. B) In the right, DCA contact prediction of the RF00167 RFAM seed family aligned using `MAFFT` with default parameters (lower triangle) compared with the one aligned using `CARNAGE`’s (upper triangle). In the left, DCA analysis of 30 designed sequences for structure A (see above) where the upper triangle shows `CARNAGE` built MSA and the lower triangle shows `MAFFT`’s.

A second commonly used analysis is coevolutionnal (11). A coevolution signal/constraint usually reflects a direct interaction between a pair of RNA nucleotides, which usually involves physical contacts, for example, atomic interactions. We tested two cases and compared them with `MAFFT` alignment tool, our baseline. We first retrieved the Purine riboswitch (RF00167) seed alignment from RFAM composed of 133 sequences. This RNA has a typical three helices type of structure captured by our alignment method and `MAFFT`. Second, we aligned 30 sequences designed to fold in structure A (see above), but only `CARNAGE` has been able to recover its underlying structure. Coevolution contacts were extracted using the maximum likelihood DCA implemented in `pyDCA` (20).

## 5 Discussion & conclusions

Motivated by the incentive to incorporate explicitly in deep learning some of our understanding of RNA physics—in this case, the RNA folding thermodynamics— we trained a model only informed by the secondary structure adopted by random sequences. The resulting embedding is used to design targeted structures, build MSAs, and cluster sequences.

We devised a heuristic for designing RNA sequences that fold into targeted secondary structures using our approach. We showed that our method performed well on the design task, although the readout network only uses the Si-seq embeddings. Since i) only at most 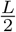 structure edges can be formed, and ii) the algorithm does not include structure predictions, the design procedure is very fast and could be used to create rapidly starting point sequences for more expensive methods.

Because trained on artificial data, Si-seq embedding is not biased by finite size datasets; therefore, unsupervised with regards to sequence similarity analysis such as the alignment and clustering. Our results showed that both the sequence and structural information are encoded in Si-seqs, at least partially, making this approach relevant for RNA functional analysis. Our strategy for the alignment and clustering task is only comparable to the state-of-the-art. Although performances could be greatly improved by optimizing the model for the alignment task (going supervised), we believe unsupervised strategies might yield a better understanding of the data. Some unsupervised approaches such as Locarna can fold and align efficiently at once, contrary to our method. However, our method has lower computational complexity and is able to handle naturally more complex structures such as pseudo knots.

RNA molecules can adopt multiple structures, so considering only the MFE structure is a real limitation. We will investigate approaches where a weighted adjacency matrix encodes multiple structures. Additionally, we will investigate strategies to incorporate more mechanistic information, such as X-Ray tri-dimensional structures and post-transcriptional modifications (epigenetics).

Graph neural networks were already applied to proteins, on the design task for example (15). By extracting the topology of graphs from the contacts observed in the X-Ray structures of known proteins, one could straightforwardly apply this approach. Although more information can be used in three dimensional structures, the training size is finite in contrast to the case of RNA secondary structures; therefore, it will require some sort of regularization.

